# Metatranscriptomic response of the wheat holobiont to decreasing soil water content

**DOI:** 10.1101/2022.09.29.510153

**Authors:** Pranav M. Pande, Hamed Azarbad, Julien Tremblay, Marc St-Arnaud, Etienne Yergeau

## Abstract

Crops associate with microorganisms that help their resistance to biotic. However, it is not clear how the different partners of this association react during exposure to stresses. This knowledge is needed to target the right partners when trying to adapt crops to climate change. Here, we grew wheat in the field under rainout shelters that let through 100%, 75%, 50% and 25% of the precipitation. At the peak of the growing season, we sampled plant roots and rhizosphere, and extracted and sequenced their RNA. We compared the 100% and the 25% treatments using differential abundance analysis. In the roots, most of the differentially abundant (DA) transcripts belonged to the fungi, and most were more abundant in the 25% precipitation treatment. About 10% of the DA transcripts belonged to the plant and most were less abundant in the 25% precipitation treatment. In the rhizosphere, most of the DA transcripts belonged to the bacteria and were generally more abundant in the 25% precipitation treatment. Taken together, our results show that the transcriptomic response of the wheat holobiont to decreasing precipitation levels is more intense for the fungal and bacterial partners than for the plant.

## Introduction

Drought is one of the most significant threats to crops and will become more frequent and intense with climate change [1]. Both the plant and its microbiota respond to decreasing soil water content, which affects the fitness of the plant. However, because of a lack of studies integrating plant and microorganisms, the best targets for improving crop resistance to water stress are not clear. Many *Actinobacteria* and *Proteobacteria* can improve plant tolerance to drought- or salinity-related stresses [2-5]. Fungal endophytes can also improve plant performance under abiotic stress [6-8]. Mycorrhizal fungi can improve water use efficiency and reduce drought stress in wheat [9], oat [10], and corn [11]. Interestingly, endophytic and rhizospheric microorganisms isolated from environments prone to drought tend to confer plants with a better resistance to drought [8, 12]. Many mechanisms are involved in the enhancement of plant drought tolerance by microbes. These include modulation of plant drought stress genes [13], reduction of the stress hormone ethylene levels through degradation of its precursor 1-aminocyclopropane-1-carboxylic acid (ACC) by the bacterial enzyme ACC deaminase [2, 3], stimulation of the expression of plant genes related to osmolytes and osmoprotectants by bacterial volatile organic compounds [14] and modulation of the plant epigenetics response to drought [15]. Plants also directly respond to water stress through genetic, molecular and physiological mechanisms [16].

A host and its microbiota form an holobiont, and their combined genomes is the hologenome [17]. Although the concept has been debated [18-21], it is useful in emphasising the role that microbial communities play in the host biology [for more details on these concepts, see 17]. The hologenome theory of evolution [22] considers the hologenome as one evolutionary unit, which provides an interesting framework for studying the adaptation of holobionts to stressful conditions. It implies that there are microbial-driven means by which holobionts can adapt to new environmental conditions [23-25]. The hologenome can change through 1) recruitment of new microbial partners from external sources, 2) amplification or reduction of the microbial partners already in place, and 3) horizontal gene transfer (HGT) from the external communities to the microbial partners already in place. These are coherent with the mechanisms of ecological community change put forward in the theory of ecological communities [26, 27], namely 1) migration, 2) selection, 3) speciation and 4) drift. At the transcriptomic level, the microbial response can stem from two mechanisms: 1) changes in the metagenome (by the three mechanisms listed above) and 2) changes in the gene expression of the members of the community. Although these two mechanisms cannot be disentangle using metatranscriptomics, the result will be the same: a change in the genes expressed within the holobiont. For the host, the transcriptomic response is limited to shifts in gene expression. We therefore hypothesized that most of the transcriptomic response of the wheat holobiont to decreasing soil water availability will be microbial. To test this hypothesis, we grew wheat under rainout shelters that let through 25, 50, 75 or 100% of the natural precipitation. Plant roots and rhizosphere were sampled, their RNA extracted and sequenced.

## Materials and methods

### Experimental design and sampling

Four rainfall manipulation treatments were set-up in 2016 at the Armand-Frappier Santé Biotechnologie Centre (Laval, Québec, Canada) using rain-out shelters that passively let through 25%, 50%, 75% and 100% of the natural precipitation. The rainfall exclusion treatments were performed using 2m x 2m rain-out shelters, which were covered with nine, six, three, or zero 2m x 16.7cm sheets of transparent plastic for the 25%, 50%, 75% and 100% treatments, respectively. The rain was intercepted by the plastic sheeting and guided in a gutter and downspout and collected in 20L buckets that were manually emptied when they were full. Two wheat genotypes were seeded under these shelters (drought sensitive, *Triticum aestivum* cv. AC Nass and drought tolerant, *Triticum turgidum spp. durum* cv. Strongfield), and the experiment was replicated over 6 fully randomized blocks, resulting in 48 plots (4 treatments x 2 genotypes x 6 blocks). Plots were seeded at a density of 500 seeds per m^2^ on May 18 (2016) and May 23 (2017). Seeds harvested from each of the plots were re-seeded in the exact same plot the following year. For the current manuscript, only the Strongfield cultivar was used, from which rhizosphere soil and root samples were taken on July 26, 2017. For rhizosphere sampling, a plant was randomly selected (avoiding the edge of the plots), uprooted and shaken vigorously to remove the loosely attached soil. Soil tightly adhering to roots after shaking was considered as rhizosphere soil and was collected in sterile 1.5 ml microcentrifuge tubes. After collecting the rhizosphere soil, roots were washed with distilled water, separated from the plant and stored in sterile 15 ml Falcon tubes. Collected rhizosphere soil and root samples were flash frozen in liquid nitrogen within a span of 2 minutes after uprooting the plant to maintain the RNA integrity. Tubes were stored at - 80°C until the samples were processed for RNA extraction. At sampling, we also collected a bulk soil sample from the center of each plot for soil water content measurement. We measured soil water content by weighing soils before and after drying overnight at 105°C.

### RNA extraction and sequencing

Total RNA was extracted from 2 g of rhizosphere soil using the RNeasy PowerSoil Total RNA Kit (QIAGEN, Canada) and 0.5 g roots using RNeasy Plant Mini Kit (QIAGEN, Canada). Extracted RNA was treated with DNAse (ThermoFisher, Canada) to remove the DNA prior to sequencing. The absence of DNA was confirmed by the lack of PCR amplification using 16S rRNA gene specific primers. Total RNA was sent for Illumina HiSeq4000 2 × 100 bp pair end sequencing at the Centre d’Expertise et de Services Génome Québec (Montréal, Québec). Libraries for rhizosphere samples were created using a microbial ribosome subtraction approach to capture all microbial transcripts, whereas libraries for root samples were created using a poly-dT reverse transcription approach to focus on the plant transcripts. The raw data produced in this study was deposited in the NCBI under Bioproject accession PRJNA880647.

### Bioinformatics

The metatranscriptome sequencing of the 24 root and 24 rhizosphere samples resulted in 2,639M reads resulting in 264 giga bases which were processed together through our metatranscriptomics bioinformatics pipeline [28]. Briefly, sequencing adapters were removed from each read and bases at the end of reads having a quality score less than 30 were cut off (Trimmomatic v0.32) [29] and scanned for sequencing adapters contaminants reads using DUK (http://duk.sourceforge.net/) to generate quality controlled (QC) reads. QC-passed reads from each sample were co-assembled using Megahit v1.1.2 [30] with iterative kmer sizes of 31,41,51,61,71,81 and 91 bases. Gene prediction was performed by calling genes on each assembled contig using Prodigal v2.6.2 [31]. Genes were annotated following the JGI’s guidelines [32] including the assignment of KEGG orthologs (KO). QC-passed reads were mapped (BWA mem v0.7.15) (unpublished - http://bio-bwa.sourceforge.net) against contigs to assess quality of metagenome assembly and to obtain contig abundance profiles. Alignment files in bam format were sorted by read coordinates using samtools v1.2 [33] and only properly aligned read pairs were kept for downstream steps. Each bam file (containing properly aligned paired-reads only) was analyzed for coverage of called genes and contigs using bedtools (v2.17.0) [34] using a custom bed file representing gene coordinates on each contig. Only paired reads both overlapping their contig or gene were considered for gene counts. Coverage profiles of each sample were merged to generate an abundance matrix (rows = contig, columns = samples) for which a corresponding CPM (Counts Per Million – normalized using the TMM method) (edgeR v3.10.2) [35]. Each contig was blasted (BLASTn v2.6.0+) against NCBI’s nt database (version downloaded from NCBI’s server on January 9^th^ 2019) and the best hit’s taxonomic identifier was used to assign a taxonomic lineage to the contig. Taxonomic summaries were performed using MicrobiomeUtils v0.9 (github.com/microbiomeutils). The metatranscriptome co-assembly, gene abundance, read count summaries and mapping statistics and other results generated by our bioinformatic workflow are provided in the companion online Zenodo archive (https://doi.org/10.5281/zenodo.7121038).

### Statistical analyses

All statistical analyses were performed in R version 4.1.0. [36]. Transcript differential abundance analyses between the 100% and 25% precipitation treatments were carried out using the EBTest function of the EBSeq library with a false discovery rate (FDR) of 0.05. Anovas were performed using the aov function of the stats package. All graphs were generated using ggplot2. The R project folder containing the R code used for data manipulation, statistical analyses, and tables and figure generation is available on our lab GitHub repository (https://github.com/le-labo-yergeau/MT_Holobiont_Wheat). The associated transcript abundance and annotation tables, the metadata, and the soil water content files used with the R code are available on Zenodo: https://doi.org/10.5281/zenodo.7096909.

## Results

### Soil water content (SWC)

There was a significant difference (p = 0.000367) between the mean SWC across the four treatments. The water content was highest in plots exposed to 100% of the natural precipitation and gradually decreased in plots receiving 75%, 50% and 25% of the natural precipitation (Fig. 1a). The SWC was of 11% at its lowest (in the 25% precipitation treatment) and of 23% at its highest (in the 100% treatment). The rest of our analyses focus on the two most extreme conditions, the 25% and 100% precipitation treatments.

**Figure 1:**
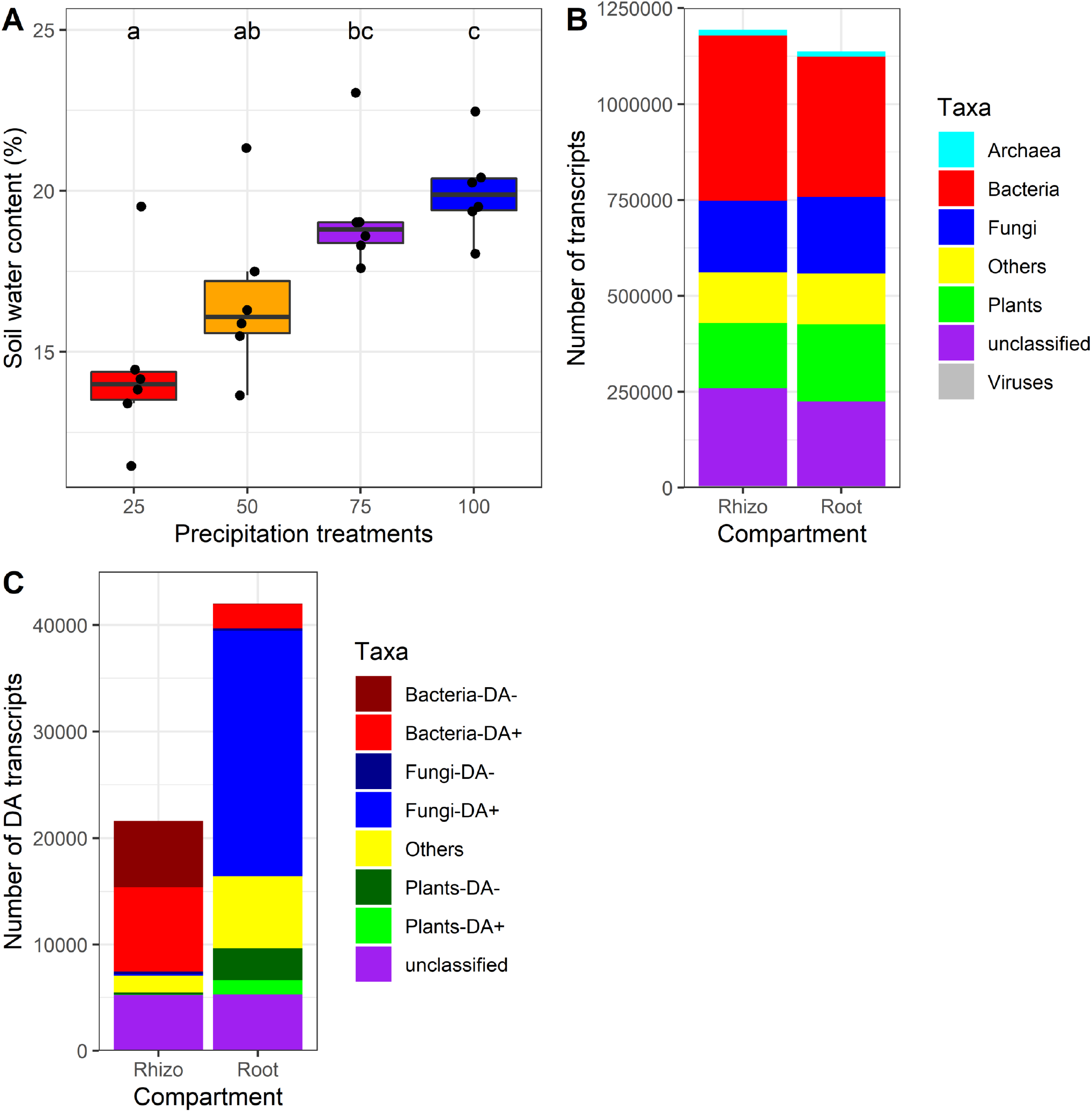
(A) mean soil water content at the time of sampling for the four different precipitation manipulation treatments, (B) kingdom-level taxonomic affiliation of the transcripts retrieved for all roots and rhizosphere samples, (C) kingdom-level affiliation of the differentially abundant (DA) transcripts together with information if they were more or less abundant in the 25% treatment as compared to the 100% treatment.

### Responses of the holobiont partners

We retrieved 1,069,108,624 clean sequencing reads (per sample, mean: 22,746,992, max: 43,912,775, min: 13,943,395) that were assembled in a total of 1,269,055 transcripts, among which 1,193,501 and 1,136,587 transcripts were found in the rhizosphere soil and wheat roots, respectively. Among the wheat root transcripts, 12,792 (1.1%) belonged to the Archaea, 365,435 (32.2%) to Bacteria, 200,233 (17.6%) to Fungi, 200,823 (17.7%) to plants, 132,313 (11.6%) to other Eukaryotes, 3,660 (0.3%) to viruses and 221,331 (19.5%) were not identified at the kingdom level (Fig 1b). Among the rhizosphere soil transcripts, 14,943 (1.3%) belonged to the archaea, 430,984 (36.1%) to the bacteria, 186,745 (15.6%) to the fungi, 169,788 to the plants (14.2%), 132,042 (11.1%) to other eukaryotes, 4,255 (0.4%) to viruses, and 254,744 (21.3%) were not classified at the kingdom level (Fig. 1b).

In the roots, among the 1,136,587 transcripts, 42,001 (3.70%) were differentially abundant (DA) at a false discovery rate (FDR) of 0.05. Among these DA transcripts, 2,309 belonged to the bacteria (5.50%), 23,274 to the fungi (55.41%), 4,357 to the plants (10.37%), 5,303 were not classified at the kingdom level (12.63%) and 6,758 belonged to other taxa (16.09%) (Fig. 1c and 2a). For bacteria and fungi, most of the DA transcripts were more abundant in the 25% treatment as compared to the 100% treatment (23,042 and 2,231 more abundant vs. 232 and 78 less abundant for fungi and bacteria, respectively), whereas it was the inverse for plant (1,295 genes more abundant vs. 3,061 less abundant) (Fig. 1c and 2a).

**Figure 2:**
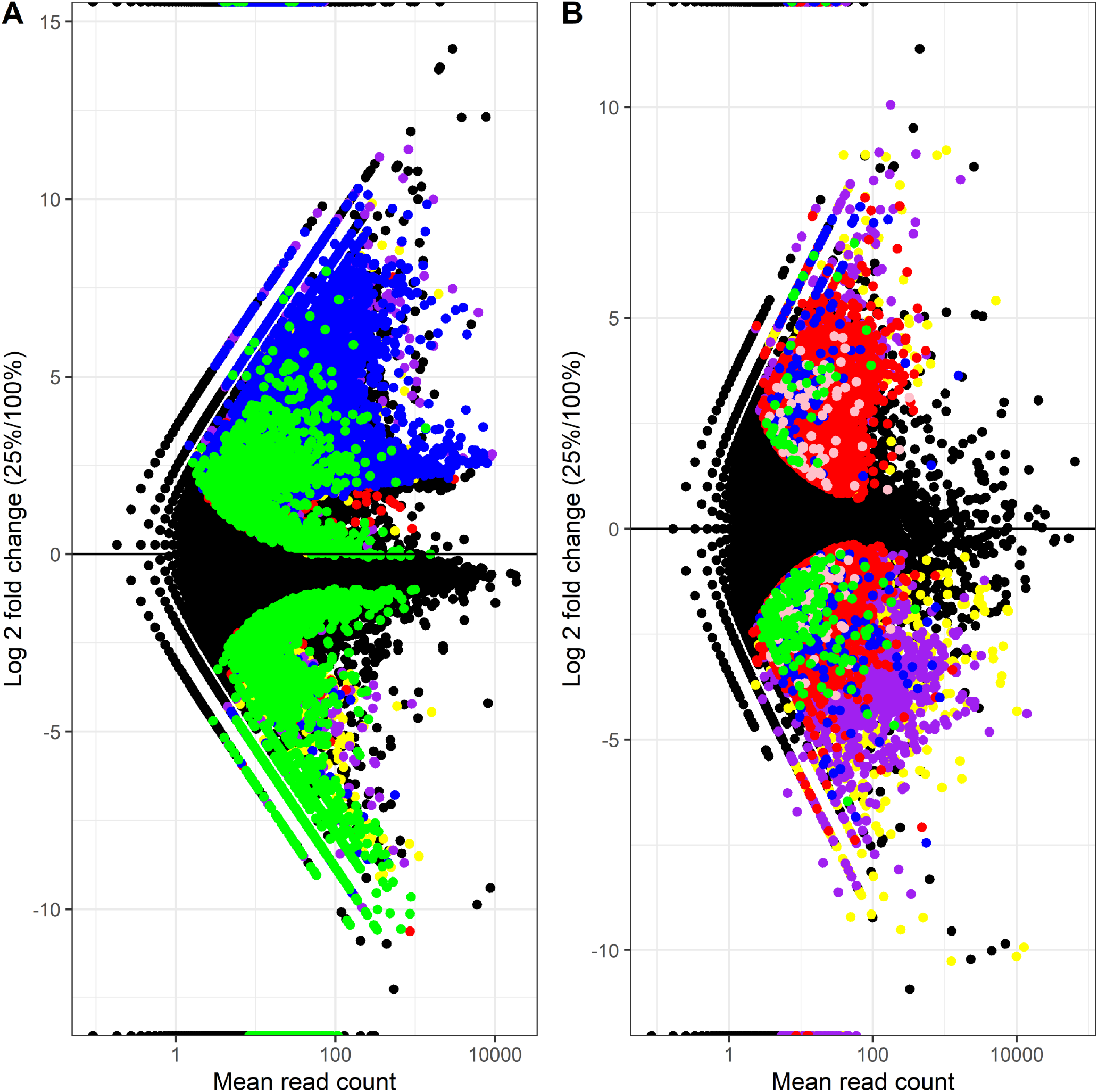
Volcano plot of transcripts log2 fold change vs. mean relative abundance, with significantly differently abundant (DA) transcripts highlighted by colors corresponding to their kingdom-level taxonomy for (A) roots and (B) rhizosphere soil. Blue: fungi, green: plant, red: bacteria, yellow: others, purple: unclassified, pink: archaea.

In the rhizosphere, among the 1,193,501 transcripts, 21,765 (1.82%) were differentially abundant at an FDR of 0.05. Among these DA transcripts, 14,178 belonged to the bacteria (65.14%), 159 to the archaea (0.73%), 402 to the fungi (1.85%), 219 to the plants (1.01%), 5,224 were not classified at the kingdom level (24.00%) and 1,583 belonged to other taxa (7.27%) (Fig. 1c and 2b). For bacteria, slightly more DA transcripts were more abundant in the 25% treatment as compared to the 100% treatment (7,938 more abundant vs. 6,240 less abundant), whereas it was the inverse for plant (41 more abundant vs. 178 less abundant) and fungi (149 more abundant vs. 253 less abundant) (Fig. 1c and 2b).

### High level taxonomy and functions of the DA transcripts

We compared the taxonomic affiliations at the phylum/class levels for all transcripts vs. for the positive and negative DA transcripts in the roots and the rhizosphere (Fig. 3). Since the DA analyses result in a single list of DA transcripts per plant compartment, we are not able to test statistically for the differences in the representation of the taxa in the different subsets. However, interesting trends emerged. Some taxa were relatively less abundant among DA transcripts than among all transcripts, suggesting a lack of response to the precipitation exclusion treatments. The Sordariomycetes, Chloroflexi, Gemmatimonadetes, among others, were in this situation across all compartments, together with the Acidobacteria in the roots and the Dothideomycetes in the rhizophere (Fig. 3). Other taxa were overrepresented among the positive DA transcripts and underrepresented among the negative DA transcripts, suggesting an increase in relative abundance or an upregulation of several genes under lower soil water content. The Actinobacteria in both compartments, the Ascomycota in the rhizosphere, and the Dothideomycetes in the roots were in that situation (Fig. 3). In contrast, some taxa were overrepresented among the negative DA transcripts and underrepresented among the positive DA transcripts, suggesting a decrease in relative abundance or a downregulation of several genes under lower soil water content. The Proteobacteria, Bacteroidetes, and Eurotiomycetes in both compartments, the Acidobacteria in the rhizosphere and the Agaricomycetes in the roots showed this pattern (Fig. 3).

**Figure 3:**
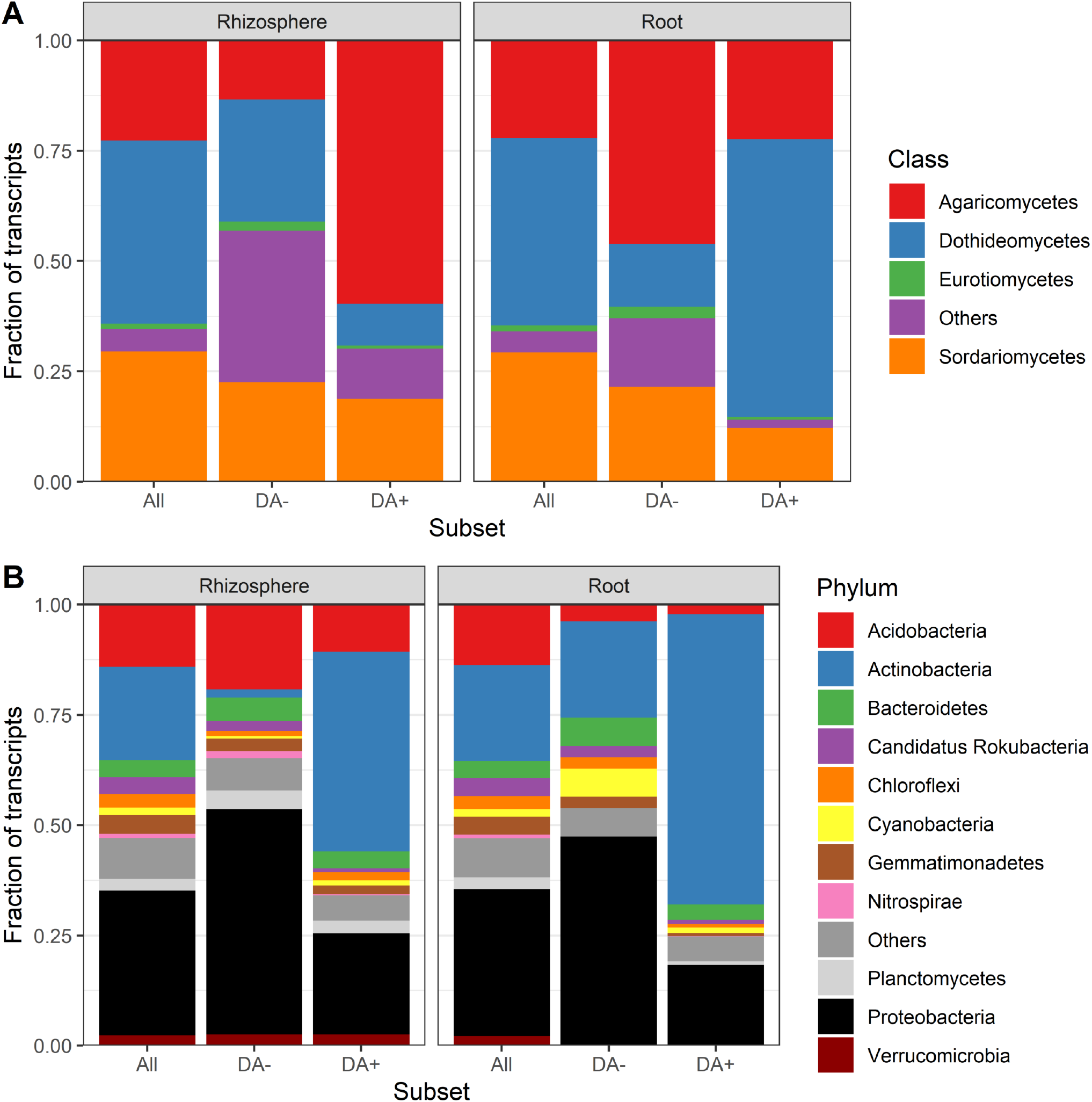
Stack bar chart comparing the taxonomical affiliations of transcripts associated to (A) fungi and to (B) bacteria in the roots and the rhizosphere across all samples vs. among transcripts positively (DA+) or negatively (DA-) differentially abundant. The transcripts not classified at this level (“NULL”) were removed.

As for COG (clusters of orthologous genes) categories, some were overrepresented in the positive DA transcripts and underrepresented in the negative DA transcripts (Fig. 4), suggesting high-level categories that are generally upregulated following a reduction of soil water content. Among these were “Carbohydrate transport and metabolism” and “Lipid metabolism” in the roots, “ “Cell envelope biogenesis, outer membrane”, “Signal transduction mechanisms”, and “Transcription” in the rhizosphere (Fig. 4). The COG categories overrepresented in the negative DA transcripts included “Translation, ribosomal structure and biogenesis” in both the rhizosphere and the roots and “Posttranslational modifications, protein turnover, chaperones” in the roots (Fig. 4). These would be COG categories that are generally downregulated with decreasing soil water content. Some COG categories were relatively less abundant among positive and negative DA transcripts than among all transcripts, suggesting a lack of response to the precipitation exclusion treatments. This included “Amino acid transport and metabolism” in the rhizosphere and “Cell envelope biogenesis, outer membrane”, “DNA replication, recombination and repair”, “Signal transduction mechanisms” and “Transcription” in the roots (Fig. 3

**Figure 4:**
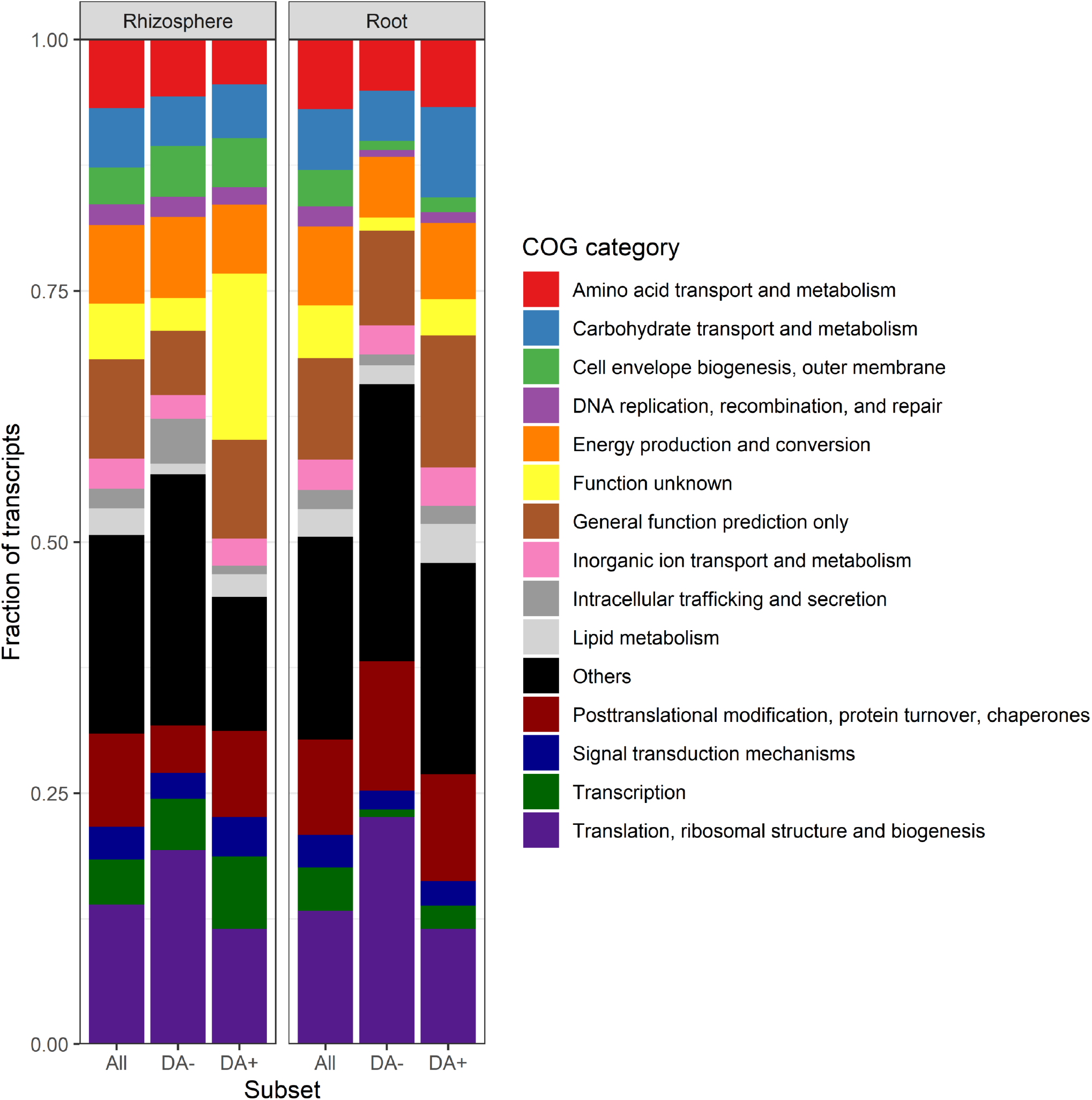
Stack bar chart comparing the functional affiliations (COG category) of transcripts associated in (A) roots and (B) the rhizosphere across all samples vs. among transcripts positively (DA+) or negatively (DA-) differentially abundant. The transcripts not classified at this level (“NULL”) were removed.

### Most differentially abundant transcripts

For DA analyses of the root samples, there were many transcripts that had a P-value=0, so we sorted them by mean abundance and are showing the top 50 transcripts in Table 1 and Figure 5A. Twenty-seven transcripts among the top 50 transcripts belonged to the *Agaricomycetes*, mostly *Coprinopsis cinerea*, and were almost all more abundant in the 25% precipitation treatment (Table 1 and Fig. 5A). Seven transcripts could be related to the wheat tribe (*Triticum aestivum* or *Aegilops tauschii*), all of which were less abundant in the 25% precipitation treatment (Table 1 and Fig. 5A). Many of the most significantly more abundant transcripts in the 25% precipitation treatment were related to amino acid and carbohydrate transport and metabolism, with transcripts such as “Amino acid transporters”, “Glycerol uptake facilitator and related permeases”, “Beta-glucanase/Beta-glucan synthetase”, “Dipeptide/tripeptide permease”, “Fucose permease”, “Neutral trehalase” and “Hexokinase” (Table 1). In contrast, many of the most significantly less abundant transcripts in the 25% precipitation treatments were linked to the COG categories “Posttranslational modification, protein turnover, chaperones” and “Secondary metabolites biosynthesis, transport and catabolism” (Table 1).

**Table 1:**
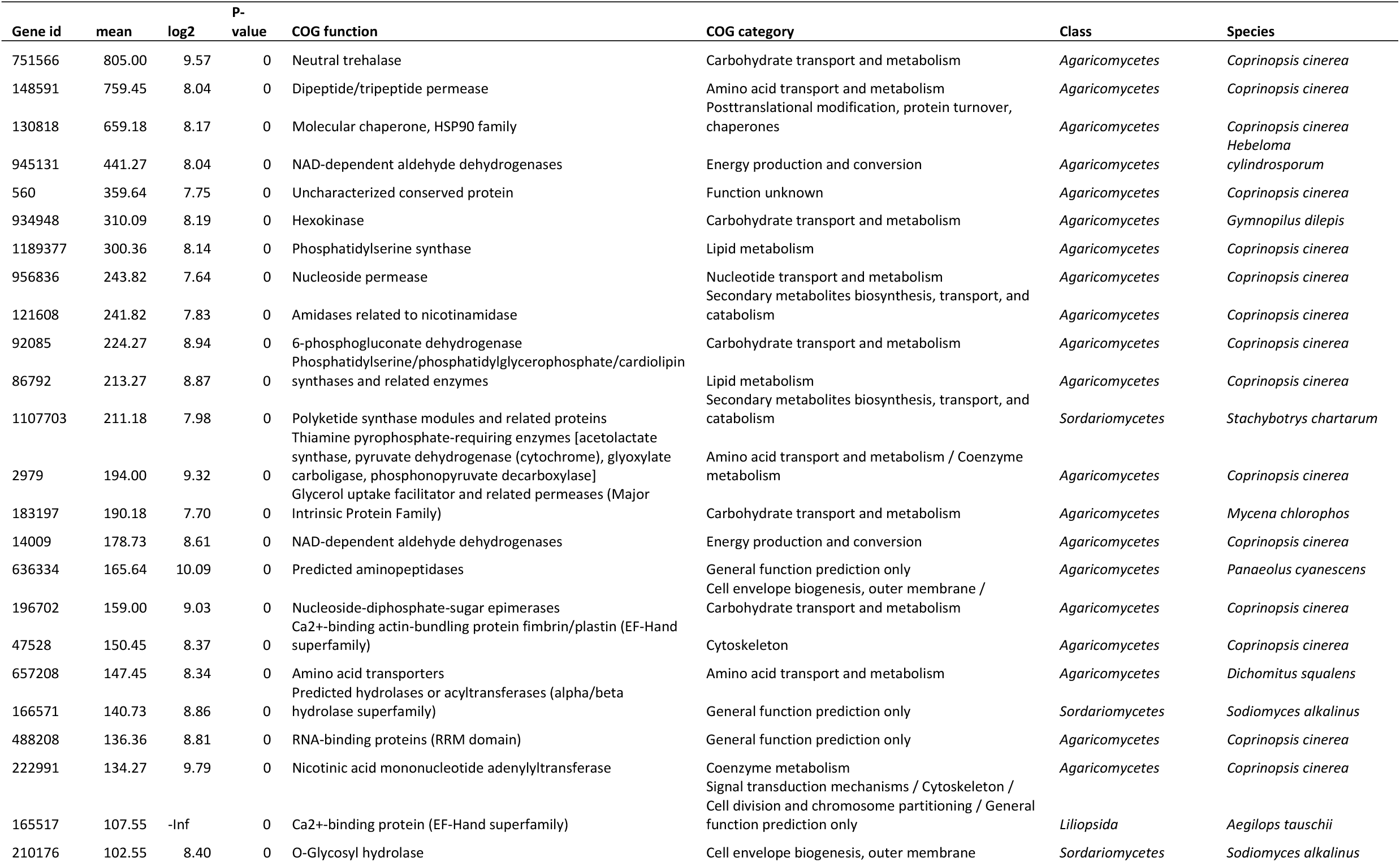
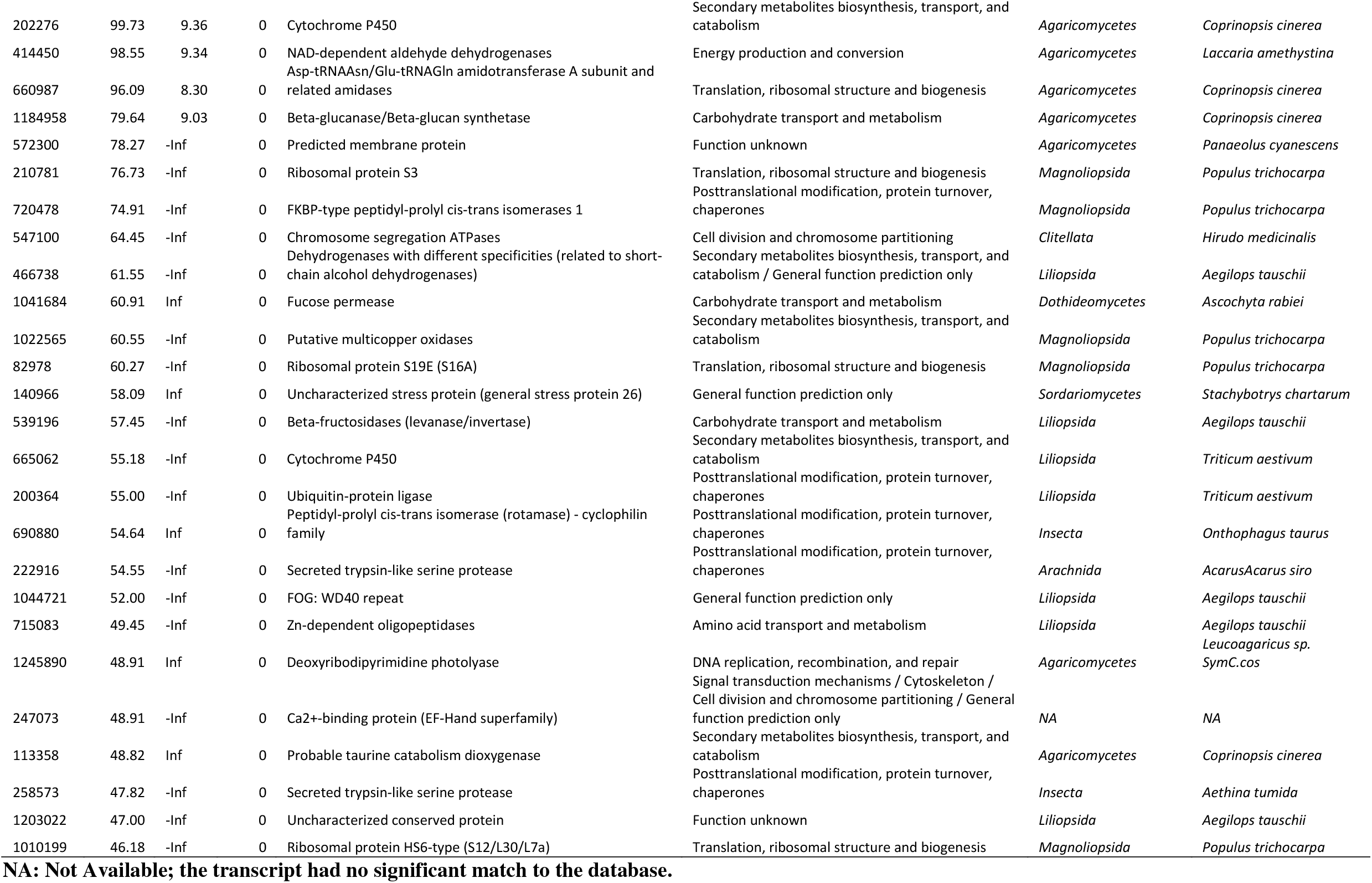
Mean abundance, log2 fold change (25% vs 100% precipitation), COG function and category and taxonomic affiliation for the top 50 most abundant genes with a P-value=0 in the roots.

**Figure 5:**
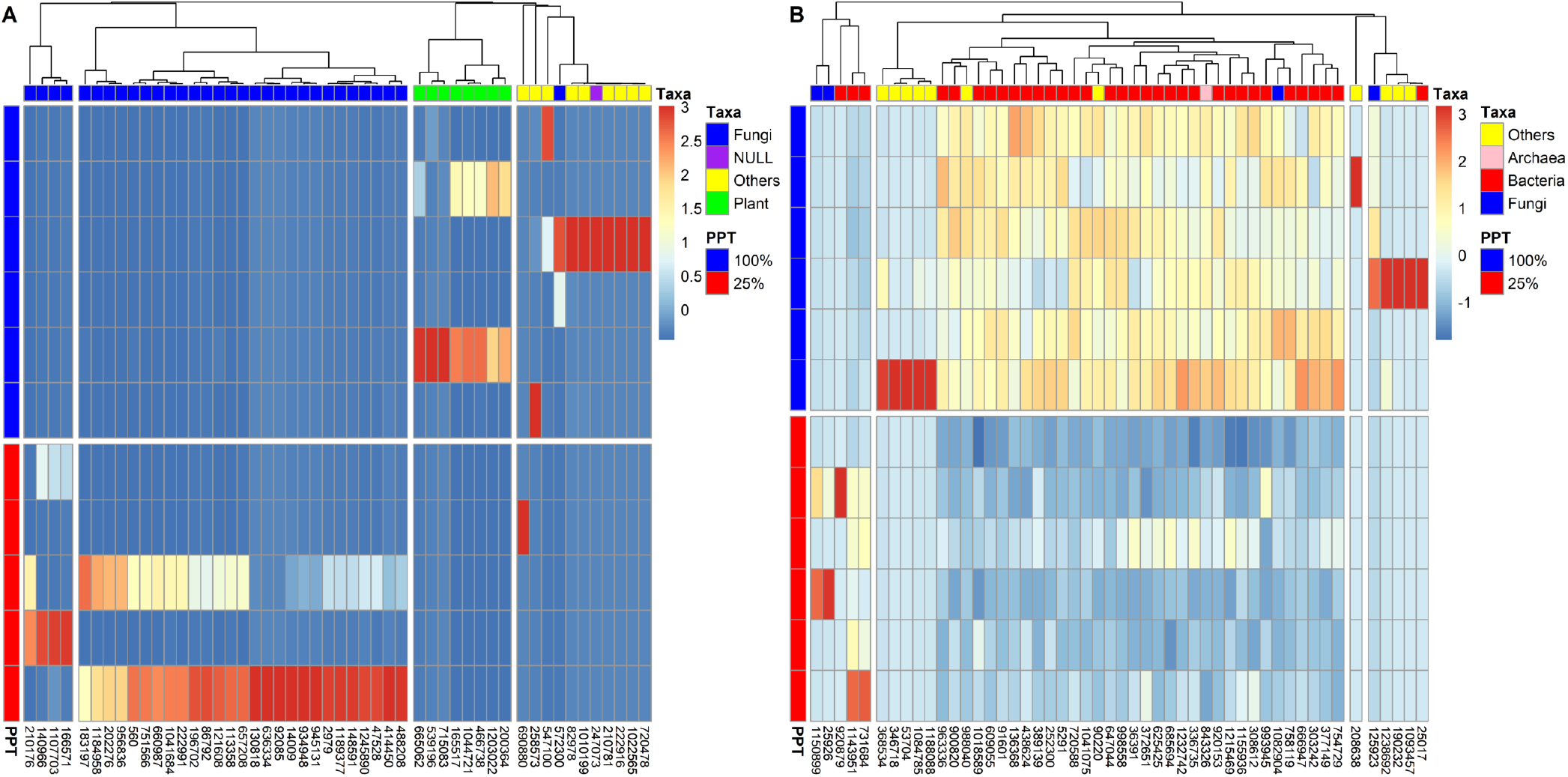
Heatmaps for the top 50 most differentially abundant transcripts for (A) roots and (B) rhizosphere samples.

For the rhizosphere, as not that many DA transcripts had a P-value=0, we are presenting the 50 lowest P-values observed in Table 2 and Figure 5B. Half the DA transcripts with the lowest P-values belonged to the *Proteobacteria*, mainly the *Alpha*- and *Delta*-classes (Table 2 and Fig 5B). Many of the most significantly less abundant transcripts in the 25% precipitation treatment were linked to the COG categories “Cell motility and secretion” and “Intracellular trafficking and secretion”, with functions related to pilus, flagella and type II and VI secretion systems (Table 2). Similar to what we observed in the roots, the fungal transcripts more abundant in the 25% precipitation treatment belonged to the *Agaricomycetes* and were related to carbohydrate and amino acid transport and metabolism (e.g. “monoamine oxidase”, “hexokinase”) (Table 2).

**Table 2:**
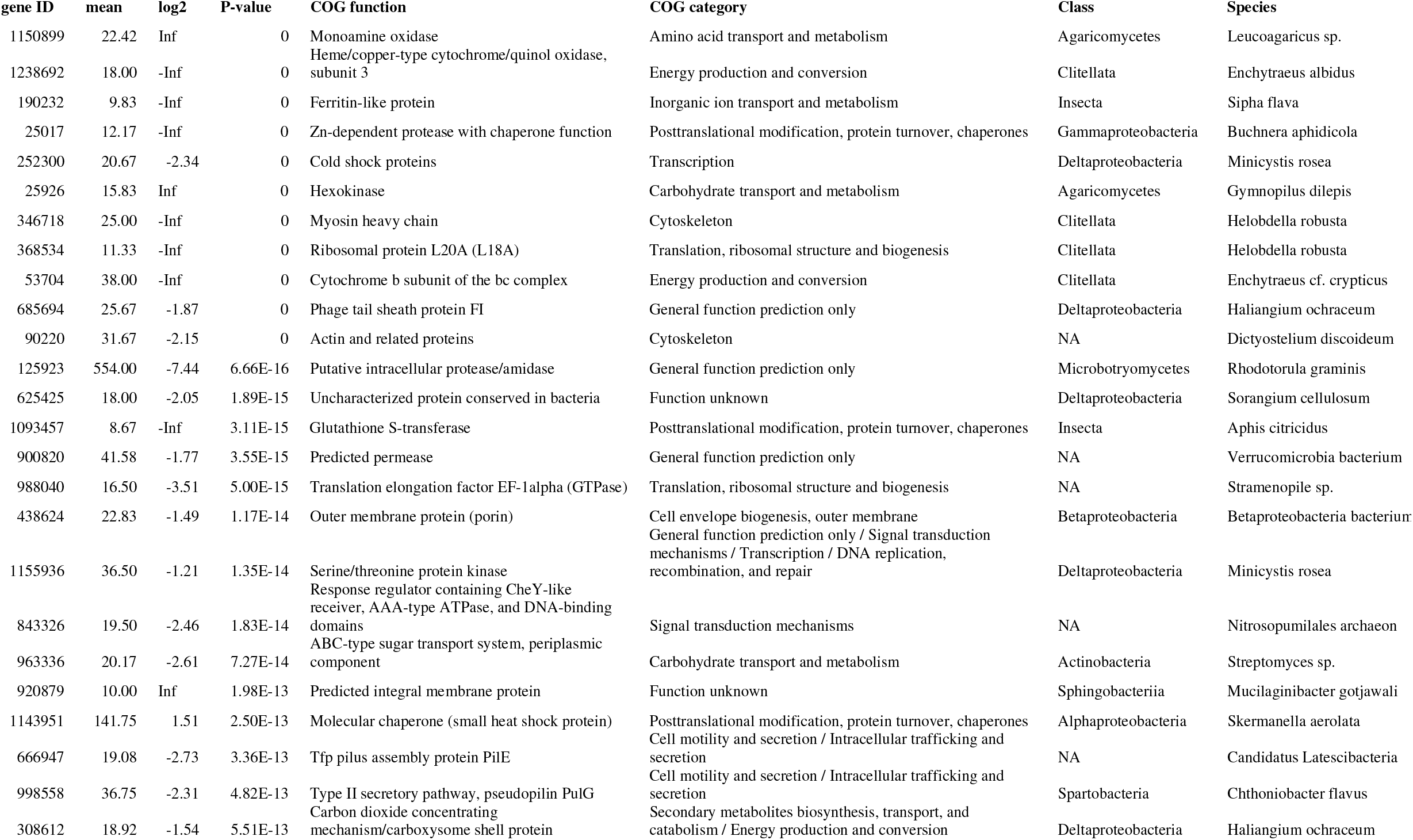
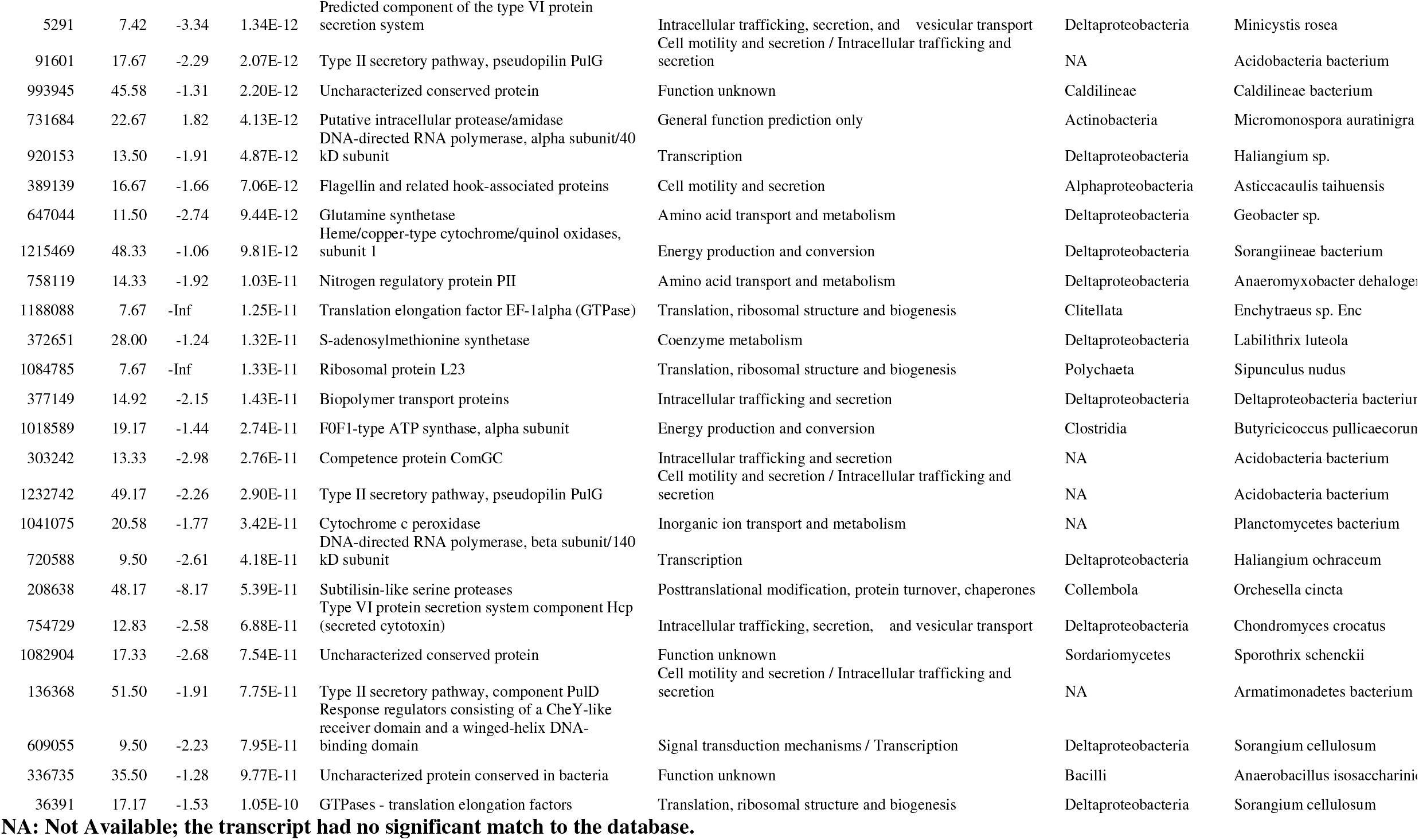
Mean abundance, log2 fold change (25% vs 100% precipitation), COG function and category and taxonomic affiliation for the top 50 microbial genes with the lowest P-values in the rhizosphere.

### DA transcripts common to roots and rhizosphere

We looked for DA transcripts that showed a common DA response in roots and the rhizosphere. Among the 37,242 and 10,565 positive DA transcripts in roots and the rhizosphere, respectively, 513 were shared (Fig. 6). Out of these 513 transcripts, 392 were affiliated to the Actinobacteria, 27 to the Basidiomycota, 12 to the Ascomycota and 11 to the Proteobacteria (Table S1). The most represented COG category were “Translation, ribosomal structure and biogenesis” (43 transcripts), “Transcription” (33 transcripts), “Carbohydrate transport and metabolism” (29 transcripts), “Posttranslational modification, protein turnover, chaperones” (26 transcripts) and “Amino acid transport and metabolism” (14 transcripts) (Table S1). Among the 4,758 and 11,200 negative DA transcripts for roots and rhizosphere, respectively, 47 transcripts were shared (Fig. 6). Most of these transcripts were not affiliated at the phylum level (26 transcripts), followed by transcripts affiliated to Streptophyta (7 transcripts) and Basidiomycota (3 transcripts) (Table S2). For COG categories, again, most of the transcripts were not affiliated with a category, and the rest were mostly affiliated to “Cytoskeleton” (5 transcripts), “Energy production and conversion” (2 transcripts), and “Translation, ribosomal structure and biogenesis” (2 transcripts) (Table S2).

**Figure 6:**
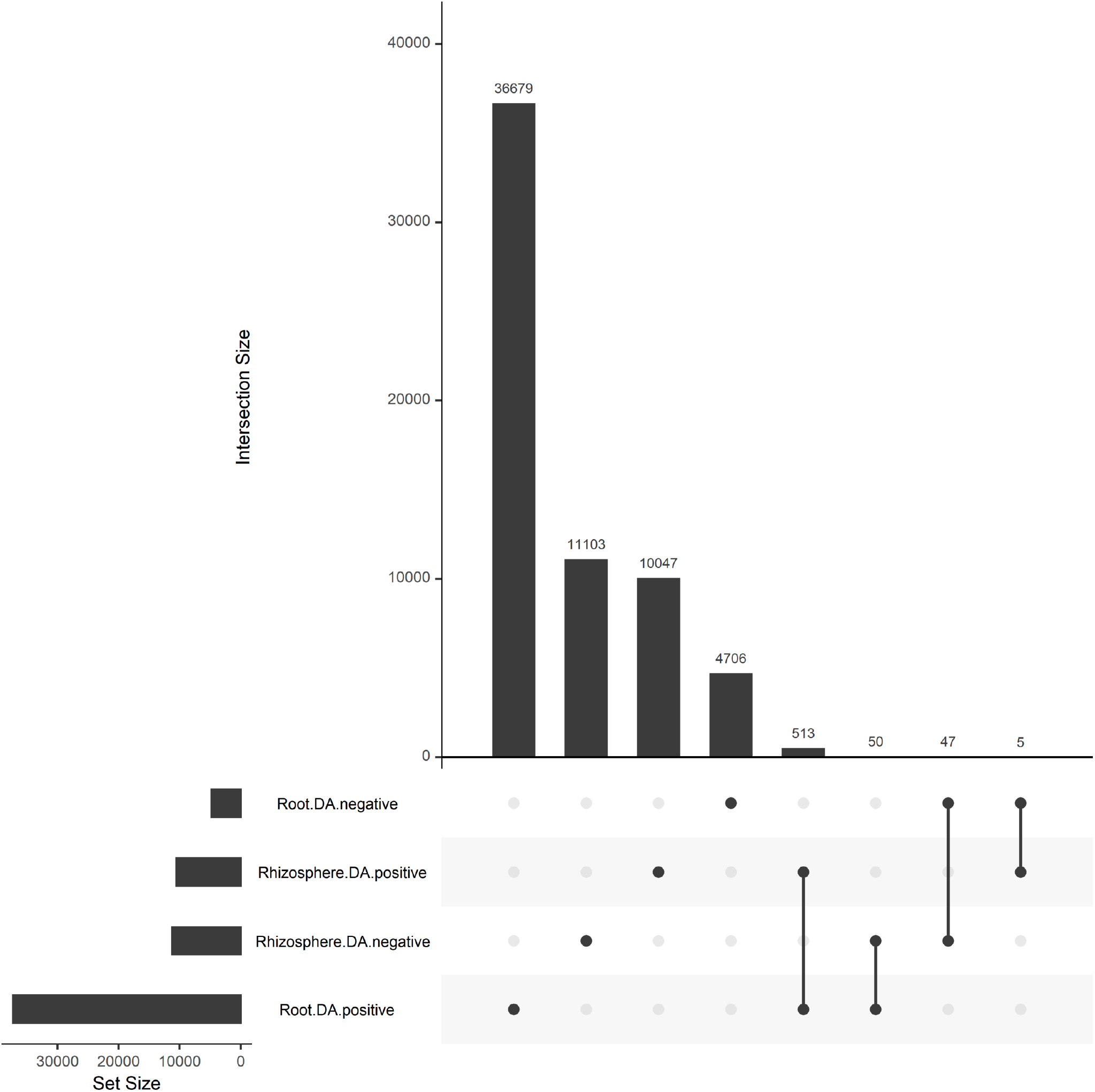
Upset plot showing the shared and unique genes between the root and rhizosphere transcripts positively (DA+) or negatively (DA-) differentially abundant.

## Discussion

We wanted to know how the wheat holobiont would respond to change in soil water availability at the transcriptomic level, and which of the partners would be more responsive. We were successful in reducing soil water content in a field experiment using rainout shelters, and found that, when comparing the two most contrasting treatments, most of the differentially abundant (DA) genes were linked to the fungi in the roots and to the bacteria in the rhizosphere. In the roots, most of the DA fungal transcripts were more abundant, whereas about half the DA bacterial transcripts in the rhizosphere were more abundant and the other half less abundant. These DA transcripts belonged to specific taxa and many of them could be related to genes known to help plants and microorganisms cope with water stress. Our results agree with one of our previous studies of the willow holobiont that showed that the root fungi are the strongest responders to soil contamination [37]. Bacteria responded mainly by expressing pollutant degradation genes, whereas plants did not show large transcriptomic responses [37]. Plant gene expression in the roots was more variable across plant genotypes than between contaminated and non-contaminated soils, in contrast to the strong response of bacteria to soil contamination [38].

The microbial component of the hologenome (the metagenome) is much more dynamic and plastic than the host genome [24]. Indeed, the microbial metagenome can be modified rapidly by changing the relative abundance of the community members, by recruiting new members from the environment or through mechanisms such as horizontal gene transfer (HGT) [24]. The host genome cannot be modified in response to environmental stress within a single generation. This could explain why most of the DA transcripts were microbial, as it combines changes in microbial gene expression and in the metagenome. The response of the host is limited to changing gene expression levels. The changes detected in plant gene expression could still affect important physiological processes, including root exudation [39]. As root exudates influence the transcriptome of bacteria [40], the microbial transcriptomic response to decreasing soil water content could have been mediated by the plant. The water depletion caused by our rainout shelters did not result in extremely low soil water content (around 12% soil water content at the lowest), which did not result in any visible stress on wheat. The wheat variety used is also water stress resistant, and this could explain the lack of a strong transcriptomic response. It would be interesting to contrast our results to the transcriptomic response of sensitive plant holobionts when exposed to much more extreme stress levels.

With the method used here, it is difficult to disentangle the metatranscriptomic response due to shifts in the composition of the microbial community and in the gene expression within the same community. The *Actinobacteria* well exemplify this. There was an overrepresentation of the *Actinobacteria* among positive DA transcripts in the rhizosphere, and most of the transcripts that were positively DA in both the roots and the rhizosphere were from this phylum. This phylum increases in relative abundance when soils get drier [41-45]. Inversely, the *Proteobacteria* and *Acidobacteria* were overrepresented among the negative DA transcripts and underrepresented among the positive DA transcripts, in line with their heightened sensitivity to water stress [45, 46]. In these two cases, the shifts observed are likely a combination of shifts in the relative abundance and of gene expression. Therefore, we referred to our differential expression analysis as a transcript differential abundance analysis. Looking only at high-level functional categories, like in Figure 4, could partly solve this problem, as general trends in gene expression at this level is less likely to be influenced by shifts in community composition. Nevertheless, we argue that whatever the underlying mechanisms are, variation in the rhizosphere and root metatranscriptome complement will have functional consequences on the holobiont adaptation to stress.

Many of the most positive DA transcripts in the roots under 25% precipitation regime, were related to amino acid and carbohydrate transport and metabolism. Amino acids, such as proline, glutamine, and glycine, betaine, and carbohydrates, such as trehalose and ectoine can be used as osmolytes [47] to maintain cellular turgor and protect macromolecular structures [48]. Gram-negative bacteria produce osmolytes purely as a drought-inducible response, whereas Gram-positive bacteria tend to produce osmolytes, at least partially, on a constitutive basis [49], which could explain some of the differences in the transcriptomic response of different taxa observed here. It would be interesting to know how much this higher abundance of transcripts is beneficial to the microbes vs. the host plant. There is some evidence that microbial endophytes and rhizobacteria can increase plant osmolyte concentration [50, 51], including proline [52], and some studies have reported that microbes can exude these compounds in the plant environment [53, 54], enabling them to directly contribute to the plant osmolyte concentration during water stress. For instance, *Coprinopsis* were often reported as endophytes of plants, including *Arabidopsis* [55] and were found here among the root fungi that showed the strongest response to decreasing soil water content, with many of their more abundant transcripts related to carbohydrate or amino acid transport and metabolism.

Other important transcripts were affected by the precipitation treatments. Among the rhizosphere bacteria, transcripts related to pilus and flagella formation were less abundant with decreasing soil water content, which might be indicative of a switch from a free-living to a biofilm lifestyle. Biofilm formation is a well-known mechanism that bacteria use to cope with environmental stresses [56]. Transcripts related to heat shock proteins were more abundant in the rhizosphere and the roots under low water content, in line with their important roles for microbes and plants under water stress [57-59]. In both root and rhizosphere, there was an overrepresentation of genes related to translation among the negative DA transcripts. A similar down-regulation of the protein biosynthesis machinery was observed in a recent soil warming metatranscriptomic study [60]. The author suggested that the increased enzymatic activity and overall metabolism caused by warming could call for a lower energy investment in ribosomes, thus optimizing resource allocation [60]. In contrast, during soil drying, *Acidobacteria* and *Verrocumicrobia* reduced their ribosomal content, whereas the *Actinobacteria* increased it [61]. Similarly, among a general decrease in translation-related transcripts, we observed here that for positive DA transcripts found in both the roots and in the rhizosphere, translation-related transcripts affiliated to the Actinobacteria was the most represented category. This differential regulation of translation among microbial groups could explain the dominance of Actinobacteria under reduce soil water availability.

In conclusion, holobionts are posited to respond in a coordinated fashion to stressful events. In our case, the microbial partners were clearly the strongest responders to decreasing water content, being responsible for most of the DA transcripts across the wheat holobiont. We had hypothesised that this would be the case since transcriptomic shifts in the microbiome combines changes in the metagenome and in gene expression, something that is not possible for the host. These transcriptomic shifts were related to microbial genes and taxa, such as the Actinobacteria and osmolyte-related genes, that are known to be beneficial to plants under water stress. Because of their dynamic response and beneficial potential, the microbiome should be considered as central in efforts to adapt crop holobionts to water stress.

## Acknowledgments

We would like to thank all the members, past and current, of the mECO:LABS for their help in setting up and maintaining the field experiment. We are also grateful to Dr. Sara Correa-Garcia for her help in making the R code portable and for submission to GitHub. This work was funded by the Natural Sciences and Engineering Research Council of Canada (Discovery grant RGPIN-2014-05274 and Strategic grant for projects STPGP 494702 to EY). We also wish to acknowledge Compute Canada for access to the University of Waterloo’s High-Performance Computing (HPC) infrastructure (Graham system) through a resources allocation granted to EY.

## Conflict of interest

The authors declare no conflict of interest.

## Notes

### Competing Interest Statement

The authors have declared no competing interest.

### Summary of Updates

Revised according to reviewers' comments.

